## Supplementary Figures for "Spike-in enhanced phosphoproteomics uncovers synergistic signaling responses to MEK inhibition in colon cancer cells"

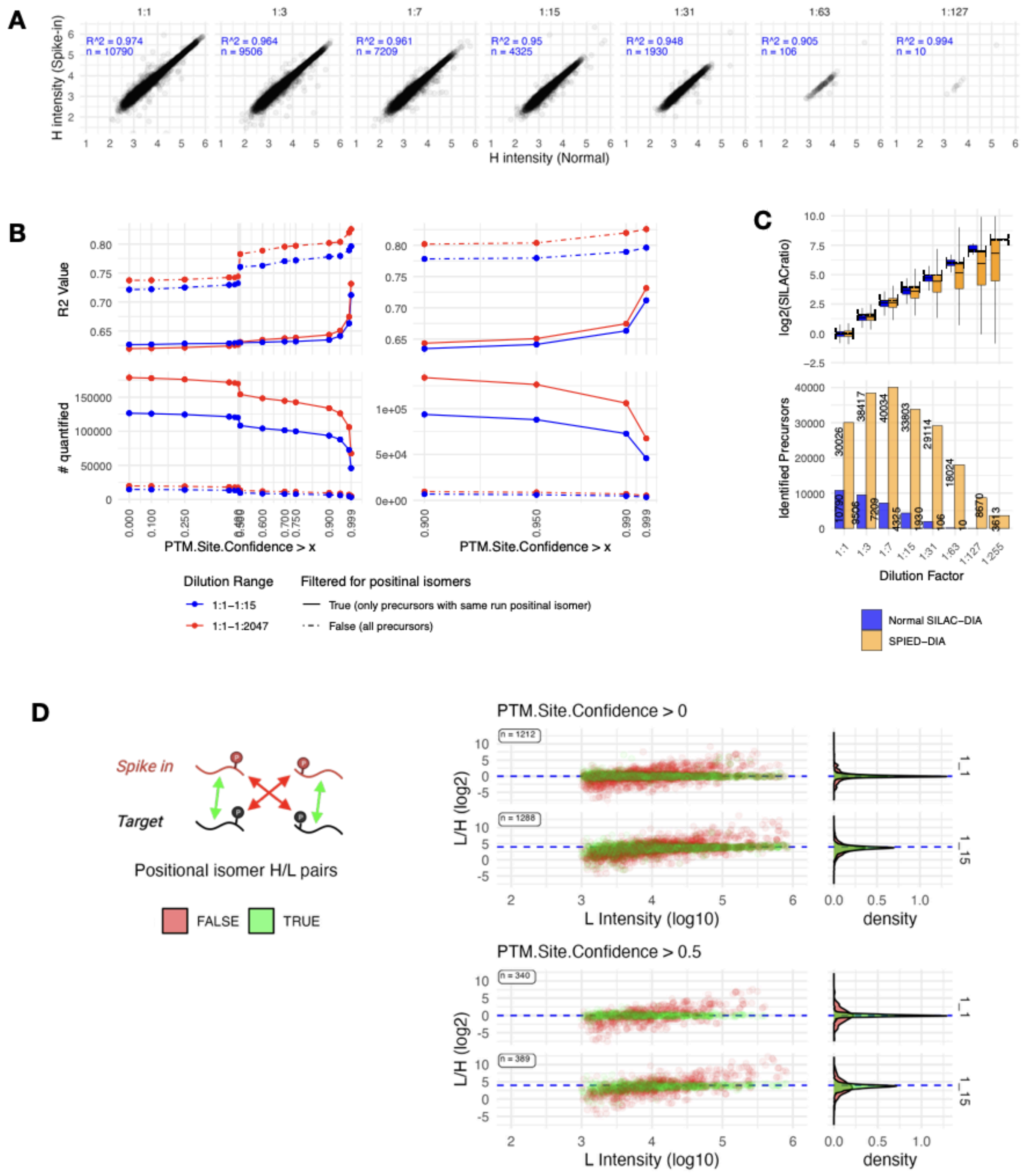


**Supplementary Figure 1: Analysis of Quantification Across Method, Dilutions and Filters.** A. Correlation of H (target) intensities between Normal SILAC-DIA and SPIED-DIA, with $R^2$ and precursor counts (n) indicated for each dilution factor. B. Effect of PTM.Site.Confidence filtering on $R^2$ values for the correlation between expected and observed SILAC ratios, contrasting first four dilutions (1:1-1:15, red) with the full range (1:1-1:2047, blue). Solid lines represent all precursors, and dotted lines represent data filtered for peptides with same run/charge positional isomers. Lower panels reflect number of precursors that survive filtering. Right panels provide a focused view on the 0.9 to 1. C. Number of identified precursors present in run dilution series without PTM.Site.Confidence filter. D. Data was filtered for peptides with same run positional isomers, and we compared H/L ratios across channel / same vs different phosphorylation site within positional isomer pairs. We looked in two dilutions how well the measured H/L ratios reflect the expected ratio within a dilution. Ratios are plotted against intensity of reference channel. Channel.Q.Value was kept consistent and PTM.Site.Confidence was varied according to the panel title

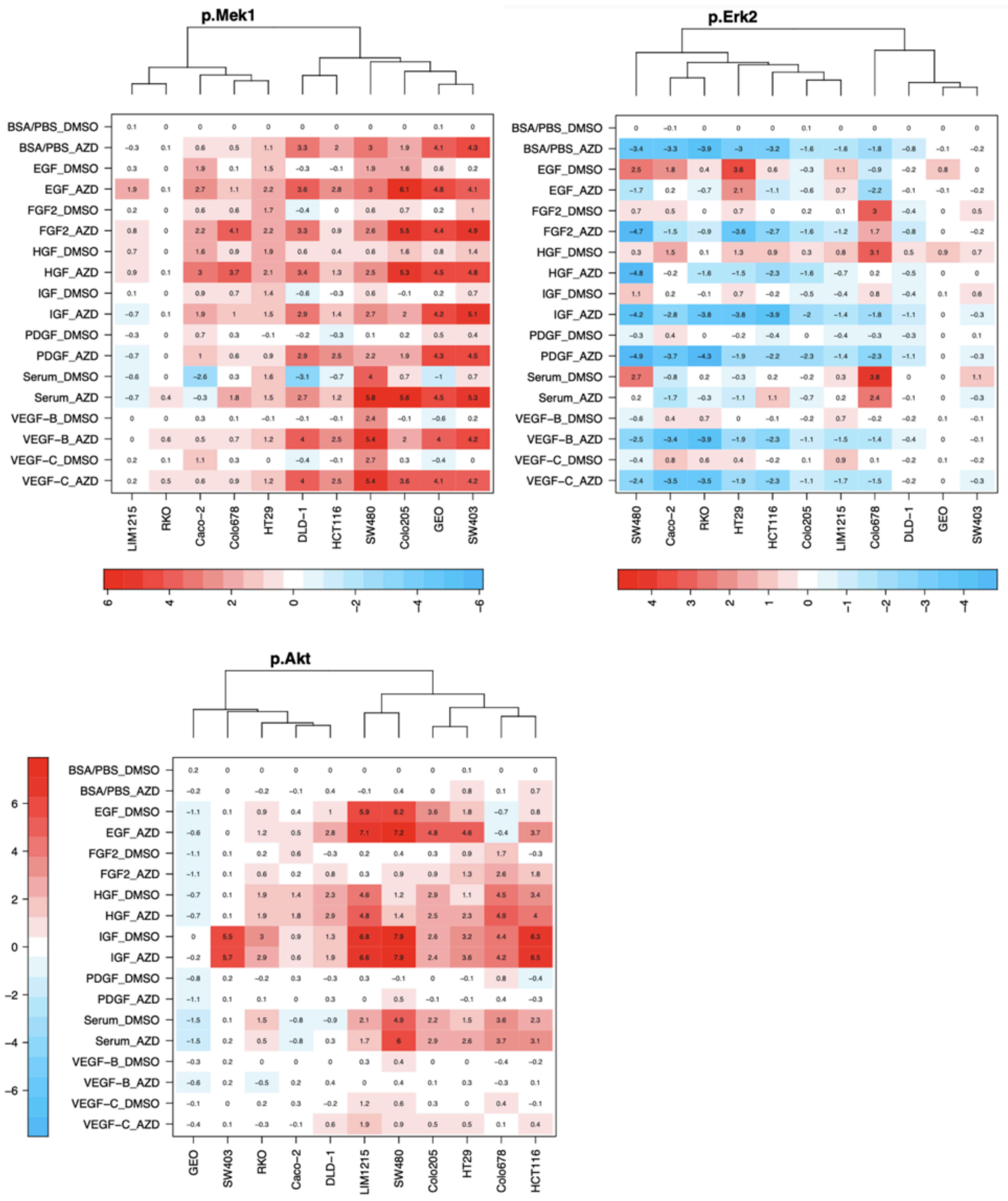


**Supplementary Figure 2: Detailed heatmaps representing luminex results from pAKT, pMEK and pERK2.** Results are separated based on MEKi ("_AZD") vs control ("_DMSO") and show intensity normalized to the BSA/PBS_DMSO condition. The expected pattern for pMEK and pERK can be observed. The cell lines per heatmap are hierarchically clustered based on response.

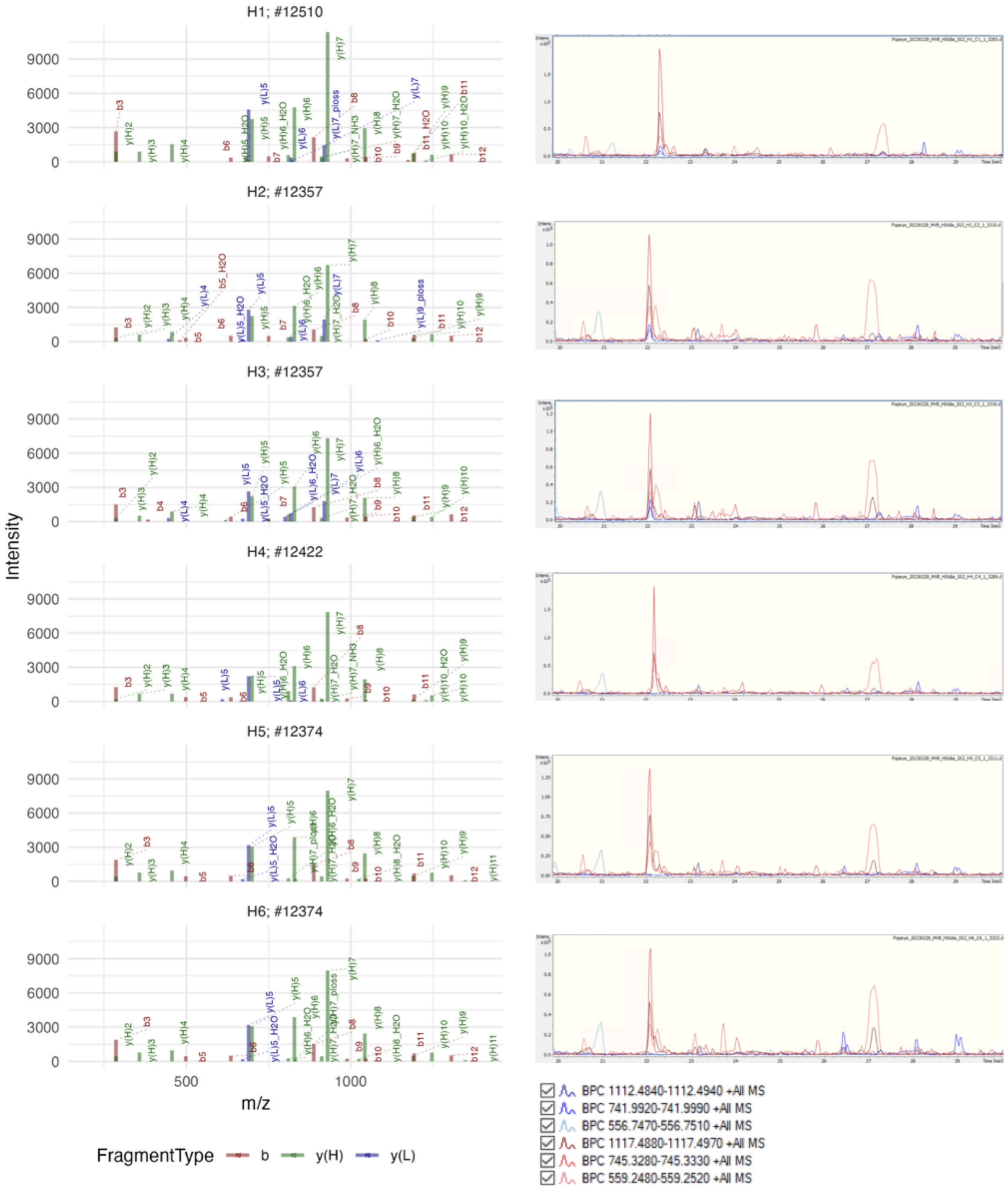
H1; #12510
H2; #12357
H3; #12357
H4; #12422
H5; #12374
H6; #12374
Intensity
9000
6000
3000
0
500
1000
m/z
FragmentType
b
y(H)
y(L)
BPC 1112.4840-1112.4940 +All MS
BPC 741.9920-741.9990 +All MS
BPC 556.7470-556.7510 +All MS
BPC 1117.4880-1117.4970 +All MS
BPC 745.3280-745.3330 +All MS
BPC 559.2480-559.2520 +All MS

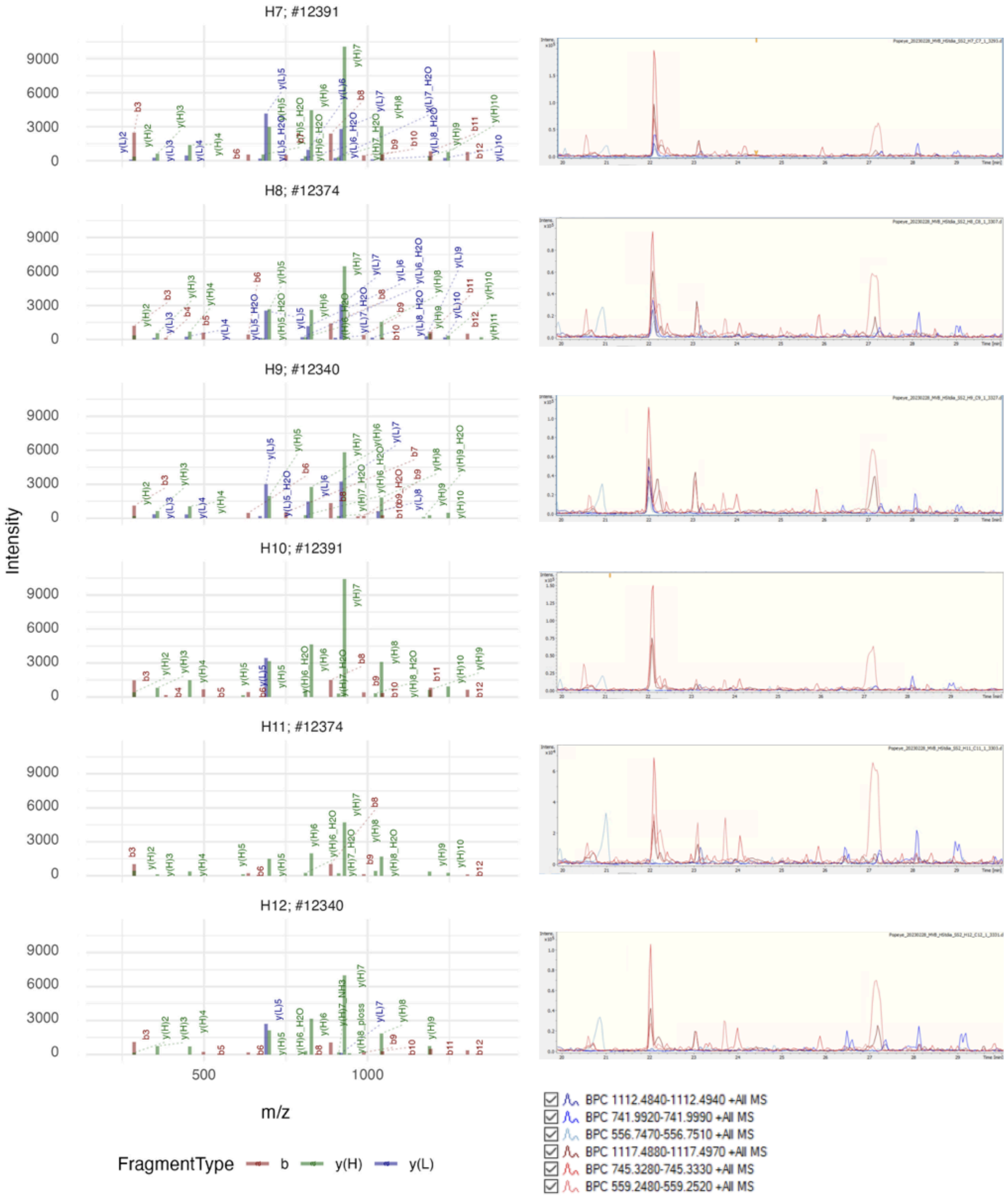


**Supplementary figure 3: Validation SILAC quantification ERK2 Tyr 187 in MS/MS spectra and MS1 traces in chromatography.** Chromatography, Spectrum and SILAC validation VADPDHDHDHTGFLTEpYVATR (3+) in HCT116 (Panel) scan number derived from DIA-NN report.tsv. Only matched ions are shown. Right hand panels show screenshot of traces of MS1 precursor masses. Blue the traces depict the endogenous precursor masses (in 2+, 3+ and 4+) and red traces depict heavy or spike in precursor masses.

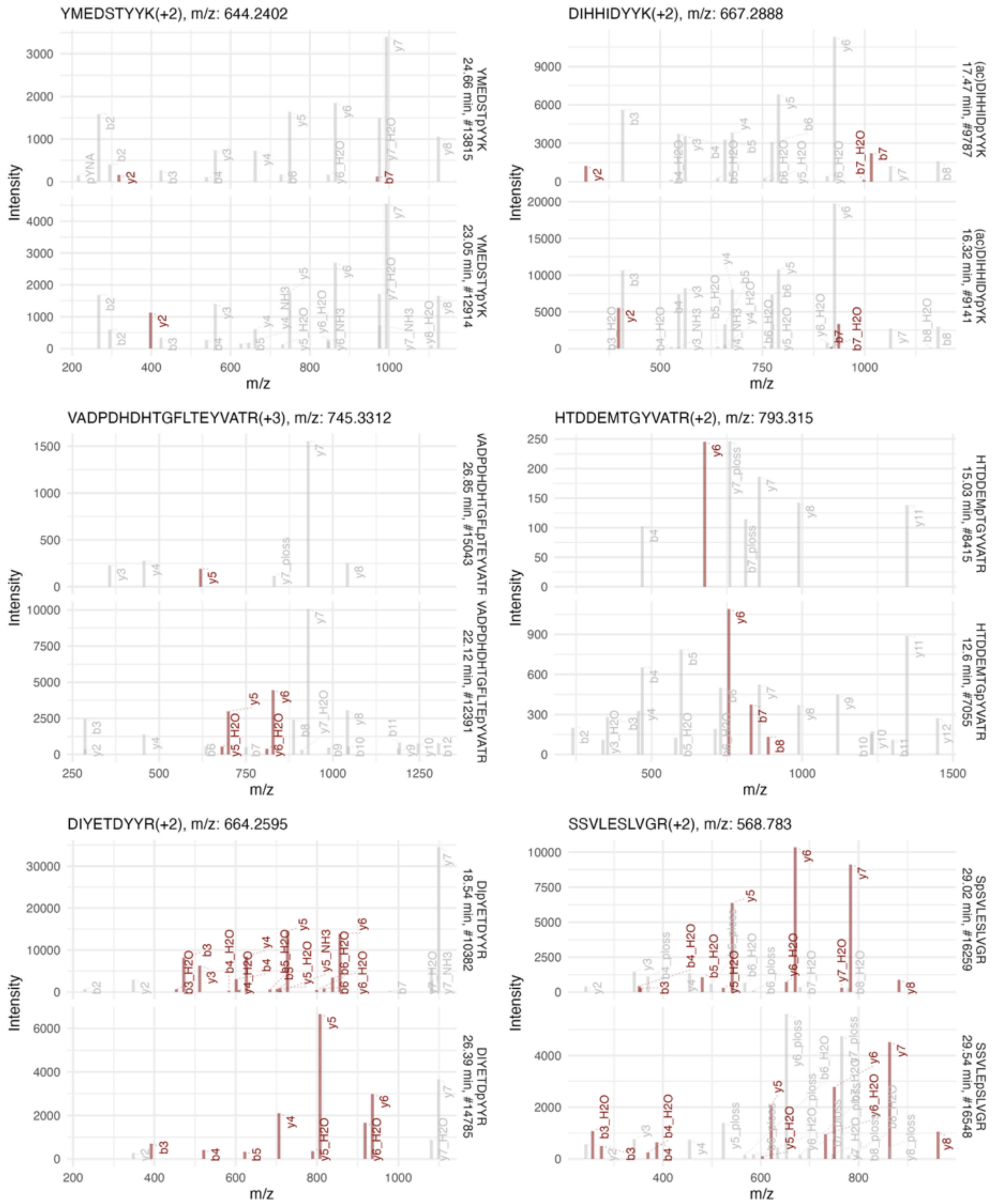

YMEDSTYYK(+2), m/z: 644.2402
YMEDSTpYYK
24.66 min, #13815
YMEDSTYpYK
23.05 min, #12914
DIHHIDYYK(+2), m/z: 667.2888
(ac)DIHHIDpYYK
17.47 min, #9787
(ac)DIHHIDYpYK
16.32 min, #9141
VADPDHDHTGFLTEYVATR(+3), m/z: 745.3312
VADPDHDHTGFLpTEYVATR
26.85 min, #15043
VADPDHDHTGFLTEpYVATR
22.12 min, #12391
HTDDEMTGYVATR(+2), m/z: 793.315
HTDDEMpTGYVATR
15.03 min, #8415
HTDDEMTGpYVATR
12.6 min, #7055
DIYETDYYR(+2), m/z: 664.2595
DIpYETDYYR
18.54 min, #10382
DIYETDpYYR
26.39 min, #14785
SSVLESLVGR(+2), m/z: 568.783
SpSVLESLVGR
29.02 min, #16259
SSVLEpSLVGR
29.54 min, #16548
Intensity
m/z

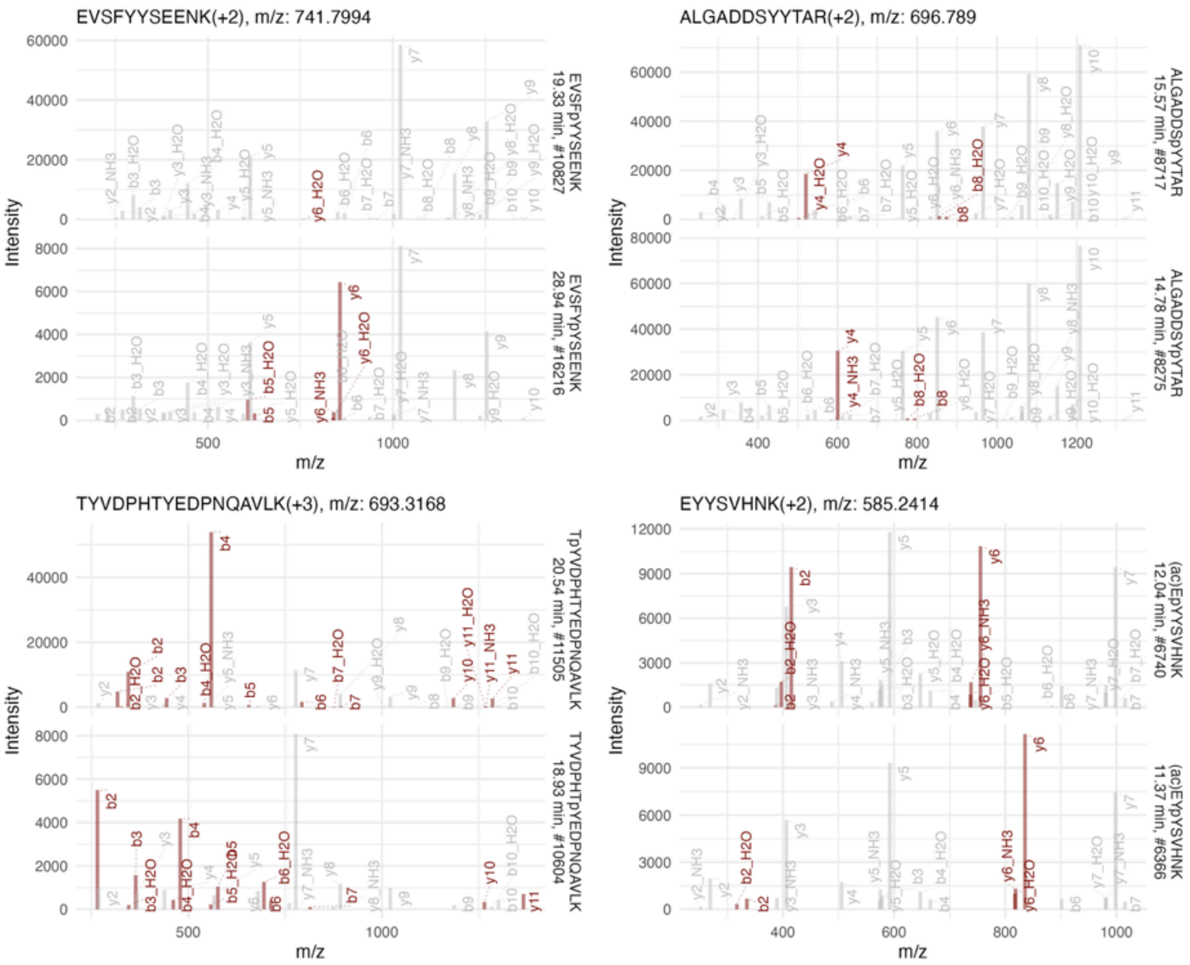


**Supplementary Figure 4: Validation of positional isomeric phosphopeptides in raw data of sample H7.** Figure represents spectra of selected phosphopeptides. We randomly selected phosphopeptides in heavy form with matching amino acid sequence and other modifications (such as n-terminal acetylation) except for phosphorylation, which is localized at different residues. Peptides are identified in identical charge states. Unmodified sequence, charge status and m/z are represented above plots. modified sequence, retention time and Bruker raw file spectrum ID are depicted at the left of plots. Fragments are labeled in accordance with fragment m/z in the library. Fragments that are crucial for distinguishing between isoforms are highlighted in red.

Note regarding Supplementary figure 4:
This highlights a particular advantage of the spike-in enabled detection. By leveraging the more intense signals from the heavy peptides, we can facilitate the identification of their light counterparts within the chromatographic run. Often, a significant retention time shift is observed between isobaric pairs, which, in combination with spectral data from the heavy peptides, allows for accurate assignment of specific phosphorylation sites, even when the relevant fragments can not be identified for the endogenous peptide.

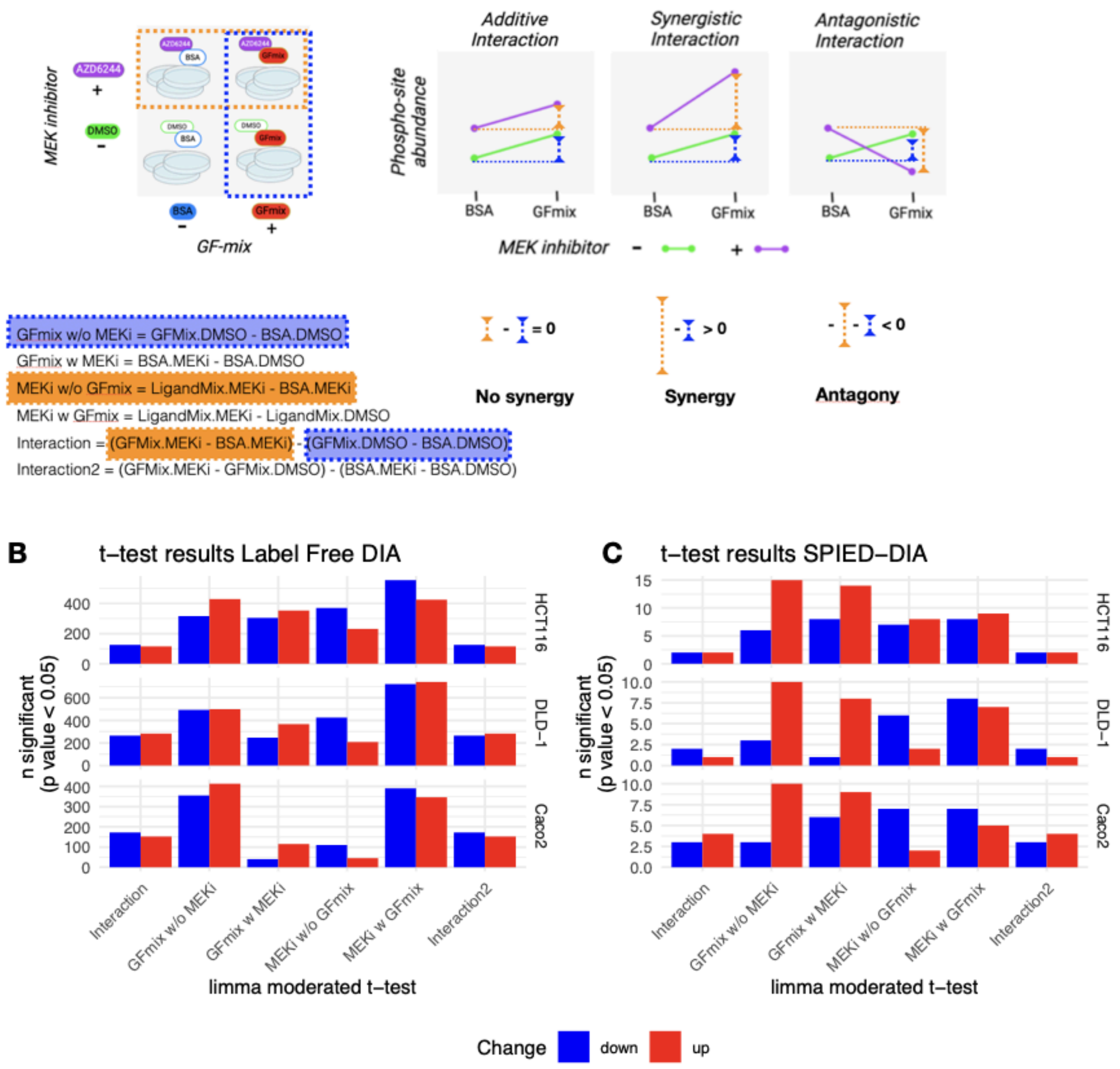


**Supplementary figure 5. Overview moderated t-test limma to test for synergistic interaction.** A. As described in Materials and methods: Within the factorial design in limma (function “makeContrasts”), contrasts were strategically defined to investigate synergistic effects: the differential impact of the growth factor mix with and without MEKi ("GFmix w MEKi" and "GFmix w/o MEKi"), and conversely, the effect of MEKi with and without the growth factor mix ("MEKi w GFmix" and "MEKi w/o GFmix"). Potential synergistic interactions were explored through an "Interaction" contrast. A linear model was fitted to the data and Bayesian statistics (“ebayes” function) were then applied to estimate variance among the precursors, employing moderated t-statistics, resulting in a logFC and (adjusted) p value per test. A positive logFC in the interaction term indicates synergistic interaction, and a negative logFC indicates antagonistic interaction. B. Results moderated t-test as defined in limma in the analysis of the label-free data. C. same as B, for the SPIED-precursors.

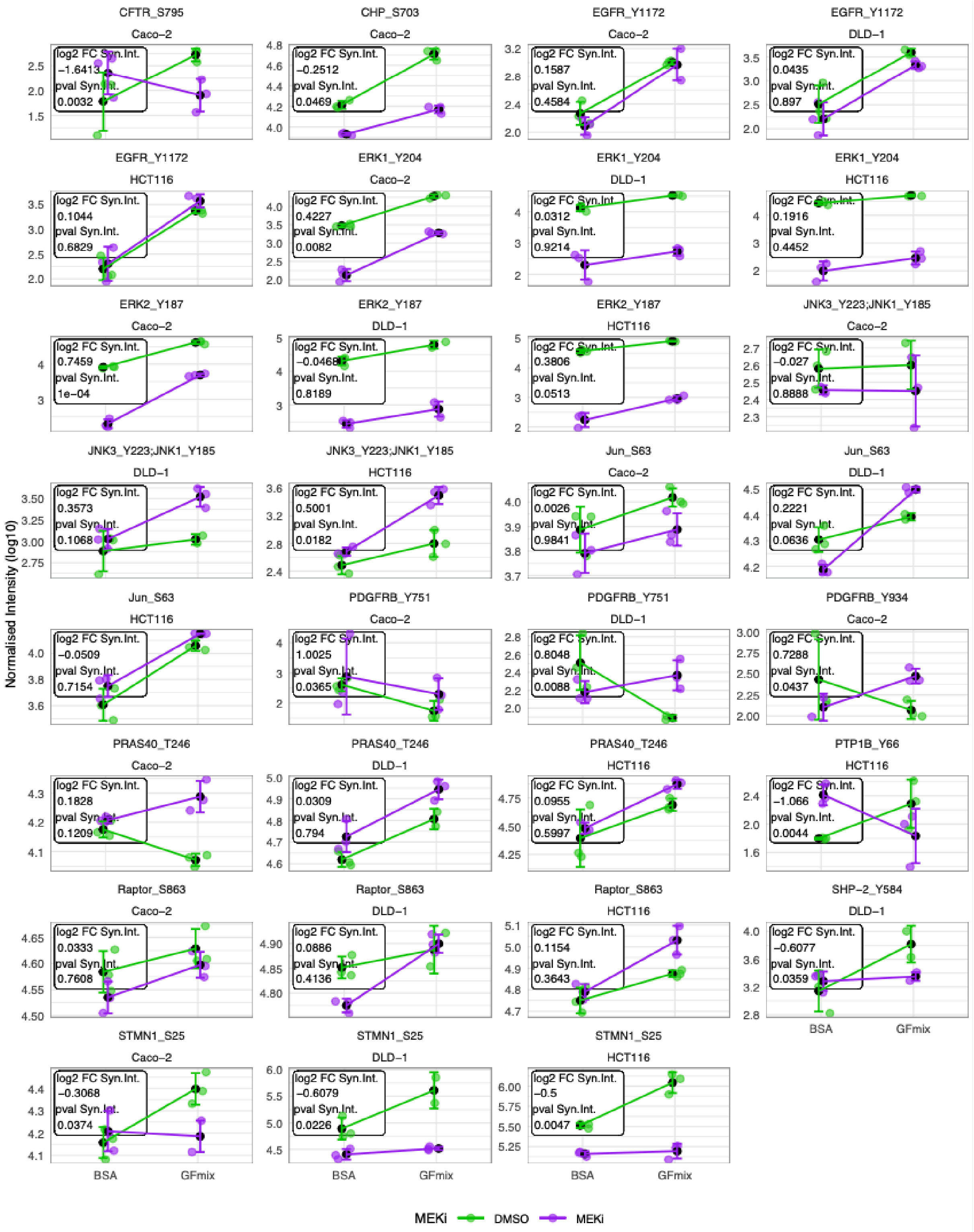


**Supplementary Figure 6. Detailed overview intensity as derived from SPIED-DIA of the phosphosites mentioned in the manuscript**. Per phosphopeptide the precursor with the lowest p-value was selected. Only precursors with at least 2 out 3 identification per condition were included in this plot.

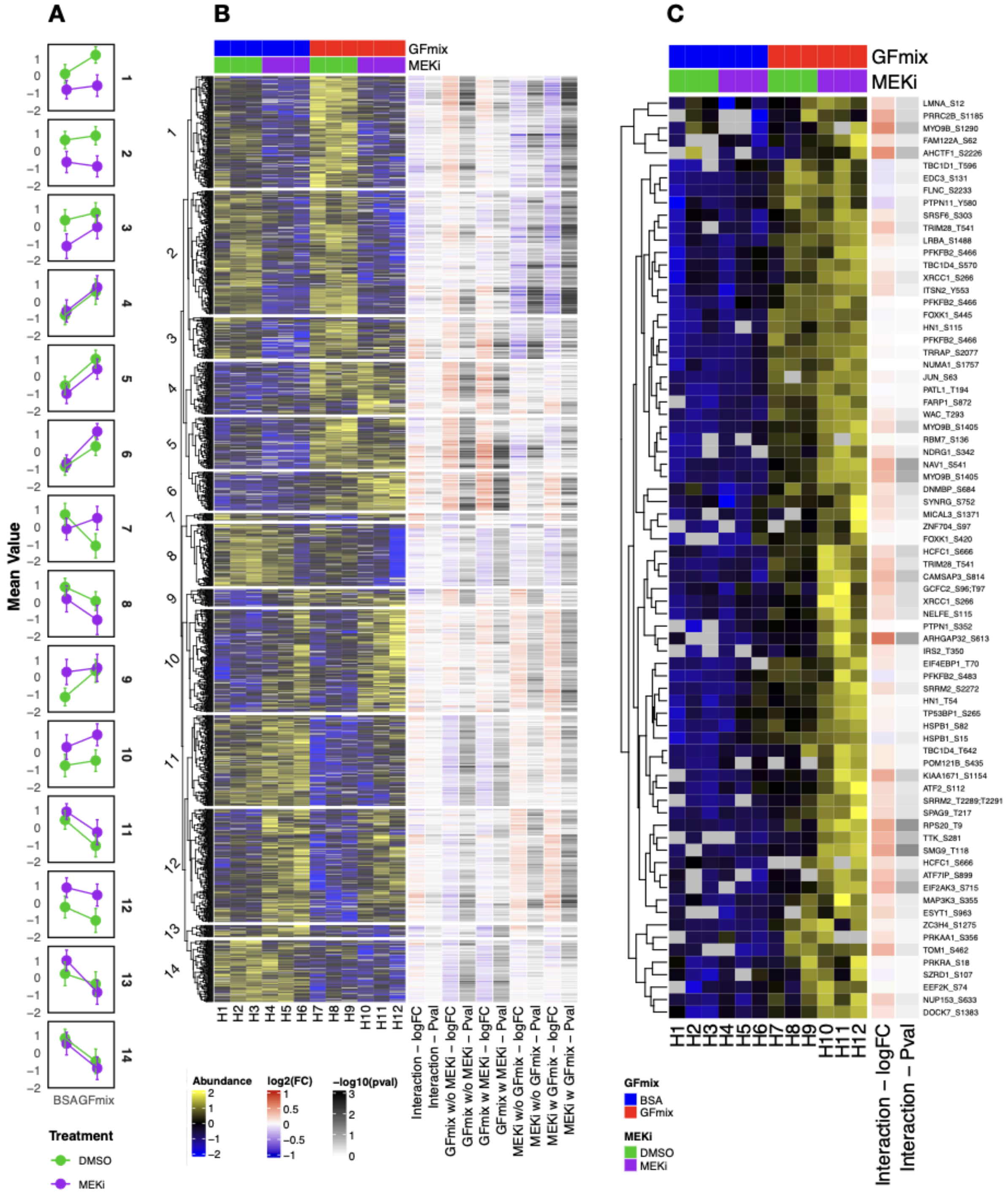

**Supplementary Figure 7: Hierarchical Clustering and Differential Abundance Analysis of phosphopeptides in label-free data HCT116** A. Mean intensity profiles per treatment condition across identified clusters. Color coding consistent with heatmap in panel B. Error bars represent standard deviation. B. Heatmap of significantly regulated phosphosites (moderated F test p-value < 0.05) from HCT116 label-free data. Row-wise z-score normalization applied to precursor intensities. Column to the right depict fold change and p-values derived from the limma moderated t-test. C. Detailed view of cluster 6. The column to the right shows log2 fold change and -log10(p-value) associated with the interaction term.

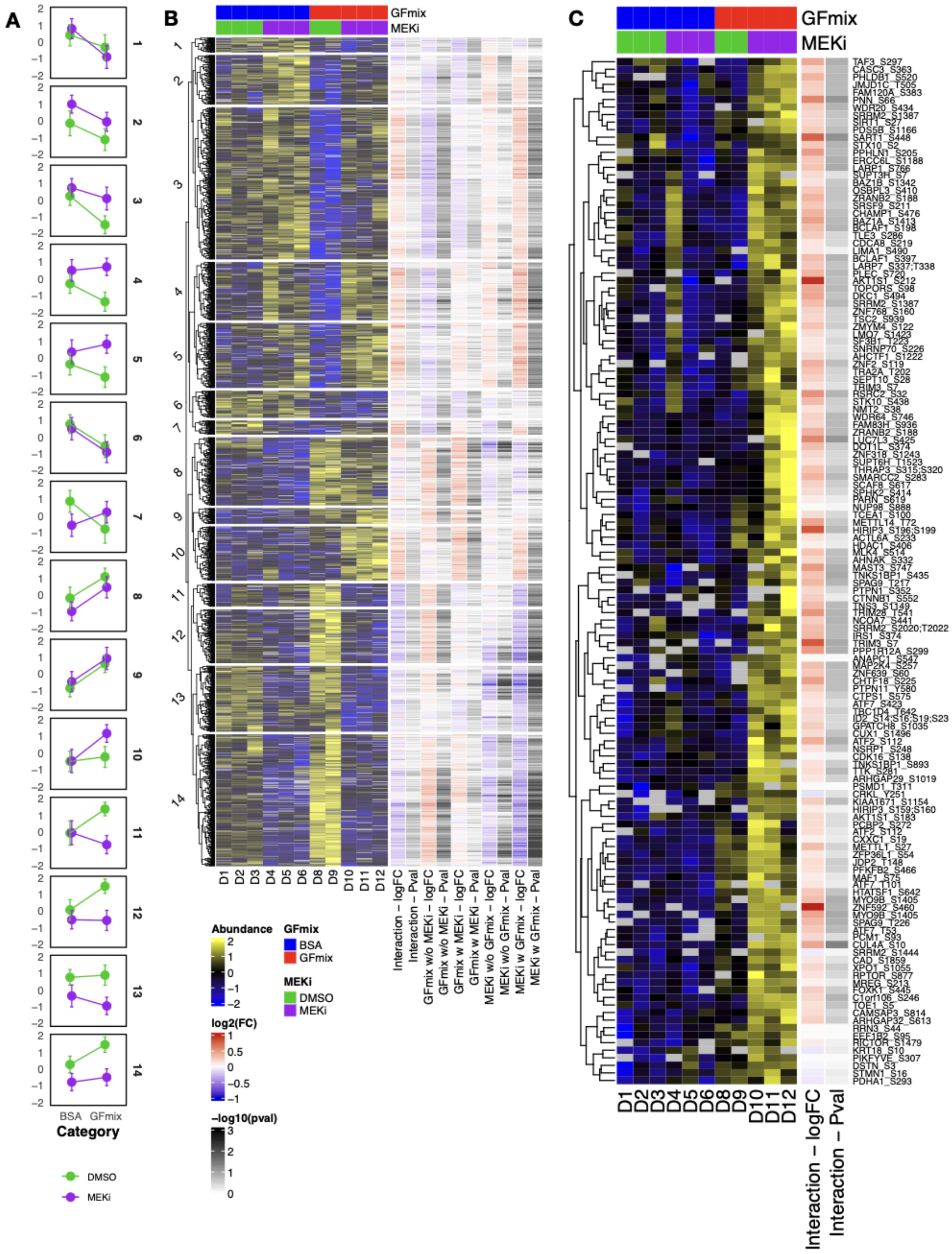


**Supplementary Figure 8: Hierarchical Clustering and Differential Abundance Analysis of phosphopeptides in label-free data DLD-1** A. Mean intensity profiles per treatment condition across identified clusters. Color coding consistent with heatmap in panel B. Error bars represent standard deviation. B. Heatmap of significantly regulated phosphosites (moderated F test p-value < 0.05) from DLD-1 label-free data. Row-wise z-score normalization applied to precursor intensities. Column to the right depict fold change and p-values derived from the limma moderated t-test. C. Detailed view of cluster 10. The columns to the right show log2 fold change and -log10(p-value) associated with the interaction term.

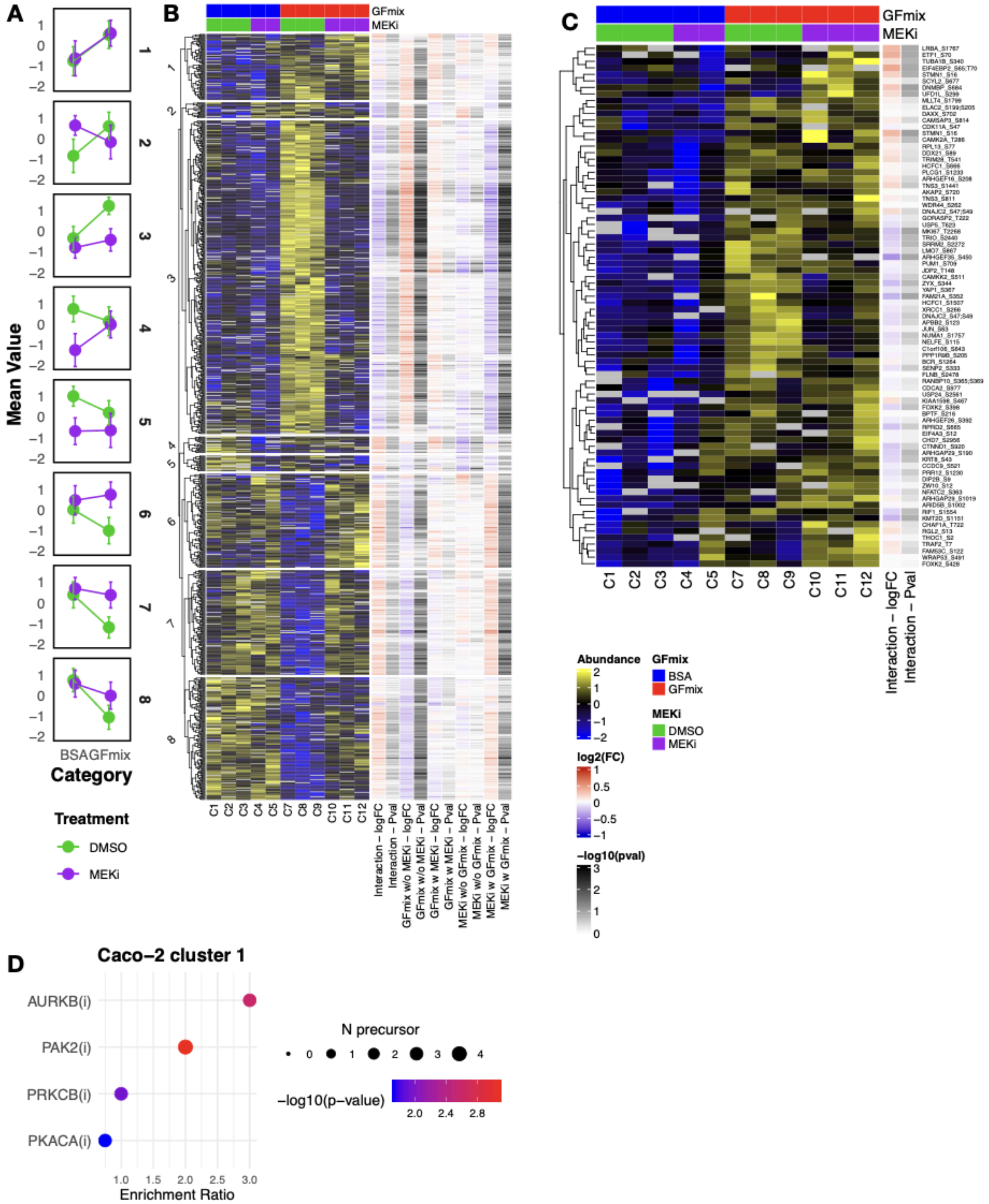


**Supplementary Figure 9: Hierarchical Clustering and Differential Abundance Analysis of phosphopeptides in label-free data CaCo2** A. Mean intensity profiles per treatment condition across identified clusters. Color coding consistent with heatmap in panel B. Error bars represent standard deviation. B. Heatmap of significantly regulated phosphosites (moderated F test p-value < 0.05) from CaCo-2 label-free data. Row-wise z-score normalization applied to precursor intensities. Column to the right depict fold change and p-values derived from the limma moderated t-test. C. Detailed view of cluster 1. The columns to the right show log2 fold change and -log10(p-value) associated with the interaction term. D. Kinase signature enrichment analysis for selected clusters from hierarchical clustered label-free data. Kinase signatures selected from PhosphoSitePlus and ikip-DB. Size and color of points indicate target count and significance as derived from Fiscers' exact test.

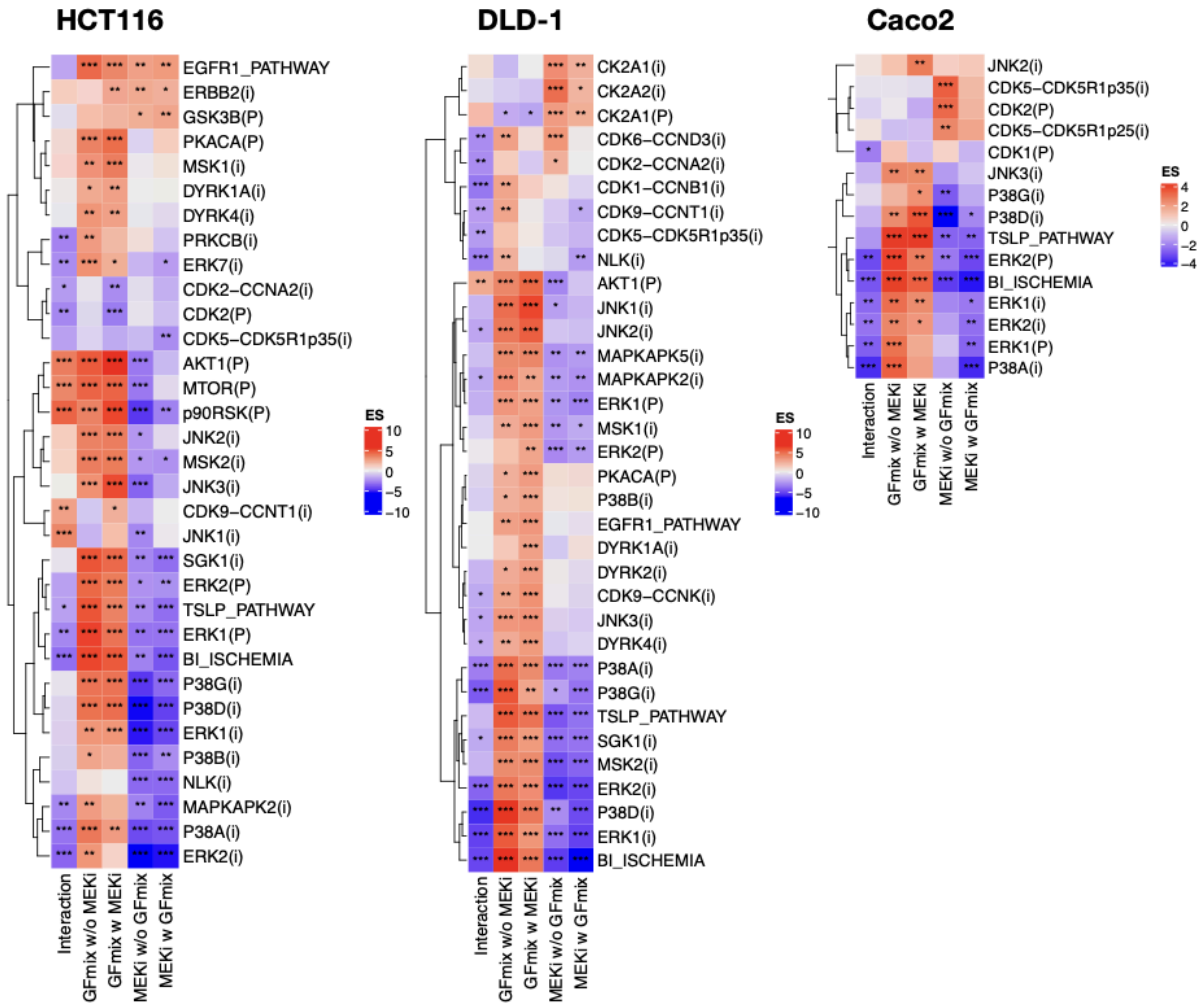


**Supplementary Figure 10. Complete results from PTM-SEA.** Signatures derived from PhosphoSitePlus (PSP) and iKiP-DB denoted by (P) and (i), respectively. The input for PTM-SEA consists of fold-change signed p-values from a moderated t-test, specifically filtered for phospho-peptides with an moderated F-test p-value <0.1, indicating significant regulation in at least one of the tests. ES = enrichment score as calculated by PTM-SEA. Significance is denoted by asterisks, with * = 0.1, ** 0.05, *** = 0.01.
